# A high-quality chromosome-scale reference genome assembly for *Asparagus racemosus* var. *CIM-Shakti* (Shatavari), a medicinal plant of Ayurvedic importance

**DOI:** 10.64898/2026.06.07.730773

**Authors:** Swati Tyagi, Ankush Sharma, Km Shivani, Vikrant Gupta, Andrew H. Paterson, Prabodh Kumer Trivedi

**Affiliations:** Plant Biotechnology Division, CSIR- Central Institute for Medicinal and Aromatic Plants, Lucknow, India; Academy of Scientific and Innovative Research (AcSIR), Ghaziabad-201001, Uttar Pradesh, India; Plant Genome Mapping Laboratory, University of Georgia, Athens, GA, 300602, USA; Department of Crop and Soil Sciences, The University of Georgia, Athens, GA, USA

## Abstract

*Asparagus racemosus* Wild., commonly known as Shatavari, is an important medicinal plant in Ayurveda and is valued for its steroidal saponins, particularly shatavarin compounds, which contribute to its adaptogenic, galactagogue, immunomodulatory, and therapeutic properties. Despite its medicinal and economic importance, genomic resources for this species have remained limited, restricting molecular breeding, pathway discovery, and comparative evolutionary studies within Asparagaceae. Here, we report a high quality chromosome scale reference genome assembly of *A. racemosus* var. CIM Shakti generated using PacBio HiFi long read sequencing and Omni C chromatin conformation scaffolding. The pseudo haploid assembly spans 817 Mb across 53 scaffolds, with a scaffold N50 of 98.50 Mb, L50 of 5, and a largest scaffold of 113.80 Mb. Ten major chromosome scale pseudomolecules were resolved, corresponding to the haploid chromosome complement of *A. racemosus*. The assembly showed high gene space completeness, with BUSCO completeness of 99.8% against the Eukaryota dataset and 98.0% against the Embryophyta dataset. BlobToolKit profiling further supported assembly quality, with GC content of approximately 39 to 40% and no major evidence of contamination. EDTA based repeat annotation identified 580.93 Mb of interspersed repetitive elements, accounting for 71.06% of the 817.57 Mb genome assembly. The repeat landscape was dominated by LTR retrotransposons, particularly Gypsy elements, which accounted for 25.01% of the assembly, followed by unclassified LTR elements at 26.58% and Copia elements at 4.84%. Structural and functional annotation identified 29,199 protein coding genes represented by 29,199 transcript models, 138,433 exons, and 125,201 CDS features. The annotation was structurally robust, with an average gene length of 4,605.1 bp, 4.74 exons per transcript, and 97.80% of transcripts containing multiple exons. The CIM Shakti reference genome provides a foundational genomic resource for investigating steroidal saponin biosynthesis, sex chromosome evolution, repeat driven genome expansion, and comparative genomics in Asparagaceae. This assembly will support future studies on medicinal trait improvement, conservation genomics, and genomics assisted breeding of climate resilient Shatavari cultivars.

## Background & Summary

*Asparagus racemosus* Willd., commonly known as Shatavari, is a dioecious perennial medicinal plant belonging to the family Asparagaceae [1]. It is widely used in Ayurveda and other traditional medicinal systems for its roots, which are rich in steroidal saponins, flavonoids, polysaccharides, and other specialized metabolites. Among these, shatavarins are considered major bioactive constituents and are associated with the plant’s adaptogenic, galactagogue, immunomodulatory, and reproductive health-related properties [2-4]. Due to increasing demand from the herbal medicine and nutraceutical sectors, Shatavari has gained substantial importance as a medicinal and aromatic plant [5]. Despite its therapeutic significance, genomic resources for *A. racemosus* have remained limited. Previous studies have relied largely on transcriptomic resources and targeted phytochemical investigations, which have provided preliminary insights into candidate genes involved in specialized metabolism [6-7]. However, the absence of a chromosome-scale reference genome has restricted systematic investigation of gene family evolution, biosynthetic pathway organization, sex chromosome biology, genome duplication history, and marker-assisted breeding. This gap is particularly important because Shatavari is a dioecious species, and chromosome-level assemblies can provide a framework for studying sex-linked regions, recombination landscapes, and structural genome evolution. In this study, we generated a high-quality chromosome-scale genome assembly for *A. racemosus* var. CIM-Shakti, an improved medicinal cultivar developed for desirable agronomic and phytochemical traits. The assembly was produced using PacBio HiFi sequencing and Omni-C scaffolding, followed by quality assessment using BUSCO, Merqury, BlobToolKit, Hi-C contact maps, repeat annotation, and structural gene prediction. The final pseudo-haploid assembly spans 817.56 Mb across 53 scaffolds, with a scaffold N50 of 98.50 Mb and 10 major chromosome-scale pseudomolecules. The assembly shows high gene-space completeness and a repeat-rich genome architecture dominated by LTR retrotransposons. The final structural annotation identified 29,199 protein-coding genes, each represented by a single transcript model. These gene models comprise 138,433 exons and 125,201 CDS features, with an average gene length of 4,605.1 bp. The high proportion of multi-exon transcripts, 97.80%, supports the quality of the annotation and indicates that the gene set is not dominated by fragmented or single-exon predictions. Functional annotation using public protein, domain, orthology, and pathway databases provides a basis for studying genes involved in steroidal saponin biosynthesis, terpenoid backbone metabolism, sterol modification, cytochrome P450-mediated oxidation, and UDP-glycosyltransferase-mediated glycosylation.

This reference genome provides an important platform for future molecular and evolutionary studies of Shatavari. It will support the identification of candidate genes underlying steroidal saponin accumulation, comparative genomics with *Asparagus officinalis* and other Asparagales species, evaluation of repeat-driven genome evolution, and development of molecular markers for breeding and conservation. The genome also provides a foundation for integrating transcriptomics, metabolomics, and population genomics to accelerate the genetic improvement of medicinal cultivars.

## Methods

### Plant material and sample preparation

Young, healthy leaves of *Asparagus racemosus* var. CIM-Shakti from a single plant were obtained from Genebank of CSIR-Central Institute for Medicinal and Aromatic Plants, Lucknow, India. The plants were grown in the genbank filed following the standard protocols. Fresh tissue was harvested in the early morning to minimize polysaccharide accumulation. High molecular weight genomic DNA was extracted using the Qiagen MagAttract HMW DNA kit following the manufacturer’s protocol with modifications to reduce polysaccharide and secondary metabolite contamination. DNA quality was assessed using pulsed-field gel electrophoresis and a Femto Pulse system, while concentration was quantified with Qubit fluorometry and Nanodrop spectrophotometry. DNA samples with a mean fragment size >50 kb and A260/280 ratio between 1.8–2.0 were used for library preparation as discussed. ^19^

### HiFi and Omni-C library preparation and sequencing

Flow-cytometric analysis [8] estimated the 2C nuclear DNA content to be 2.17 pg (Fig. 1a), corresponding to a haploid genome size of approximately 1.06 Gb. In contrast, GenomeScope analysis based on 21-mers estimated a haploid genome size of 638.4 Mb with 56.2% unique sequence content (Fig. 1b), low heterozygosity (1.1%), and low duplication levels (2.27%). The bimodal k-mer distribution indicated a diploid genome architecture with distinct heterozygous and homozygous peaks. The comparatively lower genome size estimated by GenomeScope is likely attributable to the high proportion of repetitive sequences, which are often underestimated in short-read k-mer analyses. Similar discrepancies between flow-cytometric and k-mer–based genome size estimates have been reported in several plant species, particularly in repeat-rich medicinal plant genomes. For example, Al-Qurainy et al. and Pflug et al. reported that GenomeScope substantially underestimated genome size relative to flow cytometry in *Reseda lutea, R. pentagyna* and Bembidion sp., likely due to high repeat content (57–71%) interfering with k-mer analysis [9-10]. Comparable inconsistencies between flow-cytometric and k-mer–based estimates have also been observed in other non-model plant genomes with abundant repetitive DNA. Further, SMRTbell libraries were prepared using the SMRTbell Express Template Prep Kit 3.0 (PacBio), optimized for long insert libraries. DNA repair and end-polishing steps were performed following the manufacturer’s protocol. Libraries were size-selected using a BluePippin system (Sage Science) with a cutoff of ∼15–20 kb. Sequencing was performed on the PacBio Revio platform using Revio SMRT Cells, generating high-fidelity (HiFi) reads with the latest chemistry. CCS (circular consensus sequencing) reads were produced with the PacBio CCS v6.4 pipeline, requiring a minimum of three passes per molecule and a predicted accuracy ≥0.99. In total, ∼40Gb of HiFi reads were generated. For scaffolding, an Omni-C library was prepared using the Dovetail Omni-C Kit (Dovetail Genomics) following the manufacturer’s instructions. Crosslinked chromatin was digested and proximity-ligated to generate chimeric junctions representing physical contacts across the genome. The library was sequenced on an Illumina NovaSeq 6000 platform to produce 150 bp paired-end reads. Approximately 380 million read pairs (∼150× coverage) were generated.

**Fig. 1:**
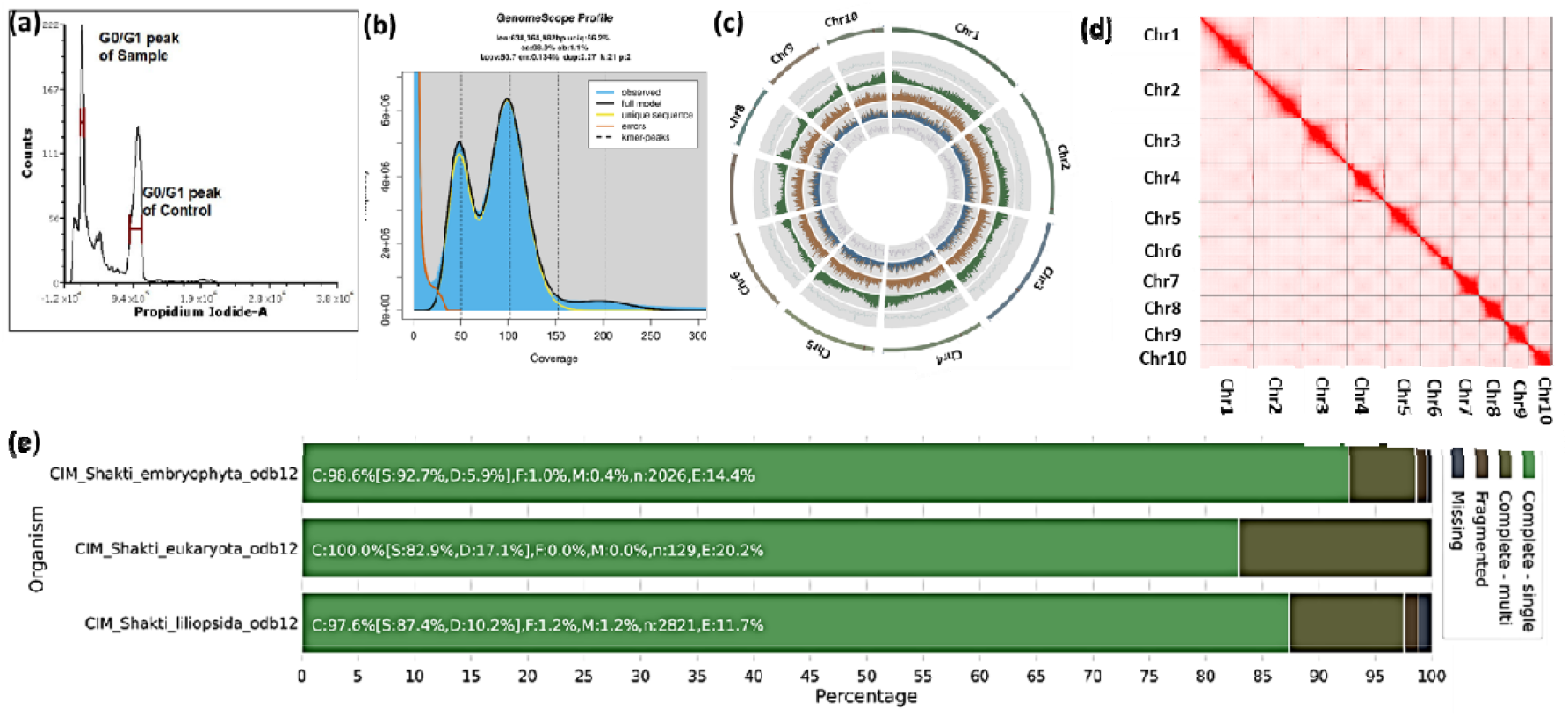
Genome characterization and chromosome-scale assembly of *Asparagus racemosus* var. CIM-Shakti. **(a)** Flow-cytometric estimation of nuclear DNA content showing the G0/G1 peaks of the sample and internal control used for genome size estimation. **(b)** GenomeScope k-mer profile (k = 21) showing the estimated genome characteristics, including genome size, heterozygosity, and repeat content based on k-mer frequency distribution. **(c)** Circos plot representation of the chromosome-scale assembly comprising ten pseudochromosomes (Chr1–Chr10). Tracks from outer to inner circles represent chromosome length, gene density, repeat density, GC content, and other genome architectural features. **(d)** Omni-C contact heatmap showing chromatin interaction frequencies among the ten pseudochromosomes, supporting chromosome-scale scaffolding and assembly continuity. Strong interaction signals along the diagonal indicate high-quality chromosome-level assembly. **(e)** BUSCO assessment of genome completeness against Embryophyta, Eukaryota, and Liliopsida datasets, demonstrating high completeness and low fragmentation of the assembled genome.

### Pseudo-haploid Genome assembly

The pseudo-haploid genome assembly of ‘CIM-Shakti’ genome was generated using Hifiasm (v.0.16.1) with default parameters optimized for HiFi reads ^11^ (Fig. 1c, Table 1). Redundant haplotigs were purged using Purge_dups (v.1.2.5) ^12^ to produce a haploid representation of the genome. Scaffolding was performed with HapHiC (v. 1.0.6) using Omni-C data, anchoring contigs into chromosome-scale scaffolds. The draft assembly comprised 53 scaffolds, totalling 817,567,924 bp, with an N50 of 98.50 Mb. These scaffolds were anchored onto 10 chromosomes (Fig. 1d), resulting a chromosome-level assembly spanned 817.57 Mb, with a scaffold N50 of 98.50 Mb, and the 10 pseudo-chromosomes together accounted for ∼98% of the genome. Assembly completeness was evaluated using BUSCO (v5.4.3) against the embryophyta, eukaryota, and liliopsida lineage datasets, which confirmed high genome integrity with 98.6%, 100.0%, and 97.6% completeness, respectively (Fig. 1e). Structural accuracy was evaluated using Merqury v1.3 which provided k-mer–based quality value (QV) estimates and base-level completeness metrics ^13^. Hi-C contact maps, visualized in Juicebox v1.11.08, confirmed the accuracy of chromosome anchoring (Fig. 1d). Haplotype-resolved assemblies were further generated from the unitig assembly obtained through *hifiasm*, following the approach described by Mascher *et al*. [14]. The final manually curated pseudomolecules for the two haplotypes spanned 921 Mb (Hap1) and 787 Mb (Hap2), respectively (Table 1). Chromosomes within each haplotype were designated according to their corresponding chromosomes in the pseudo-haploid reference. In total, 10 chromosomes were grouped for Hap1 and Hap2, providing a fully resolved, chromosome-level haplotype assembly (Fig. 2a). Both haplotypes show strong overall collinearity with the reference and with each other, with most aligned sequence falling on the expected counterpart. Overall, the haplotype-phased assembly appears largely collinear across the 10 chromosomes, with only small local rearrangement-like signals.

**Table 1.** Summary statistics of the *Asparagus racemosus* (‘CIM-Shakti’) genome assemblies.

| Metric | Pseudohaploid | Hap1 | Hap2 |
| --- | --- | --- | --- |
| Assembly Size (bp) | 817567924 | 921116963 | 787317804 |
| No. of Sequences | 53 | 1217 | 605 |
| Largest Sequence (bp) | 113801070 | 124299916 | 116002278 |
| N50 (bp) | 98,501,070 (n=4) | 95,476,940 (n=5) | 82,590,641 (n=4) |
| N60 (bp) | 93,381,493 (n=5) | 90,012,089 (n=6) | 77,057,050 (n=5) |
| N70 (bp) | 84,687,990 (n=6) | 64,956,550 (n=7) | 58,504,978 (n=7) |
| N80 (bp) | 61,906,554 (n=7) | 56,414,647 (n=8) | 53,305,368 (n=8) |
| N90 (bp) | 50,804,473 (n=9) | 53,178,899 (n=10) | 52,889,003 (n=9) |

**Table 2.** Structural annotation statistics of the *Asparagus racemosus* var. CIM-Shakti genome.

| Feature | Count / Value |
| --- | --- |
| Protein-coding genes | 29,199 |
| Transcript models | 29,199 |
| Transcripts per gene | 1.000 |
| Exons | 138,433 |
| CDS features | 125,201 |
| Exons per transcript | 4.741 |
| CDS features per transcript | 4.288 |
| Average gene length | 4,605.1 bp |
| Single-exon transcripts | 642 / 2.20% |
| Multi-exon transcripts | 28,557 / 97.80% |

**Fig. 2.**
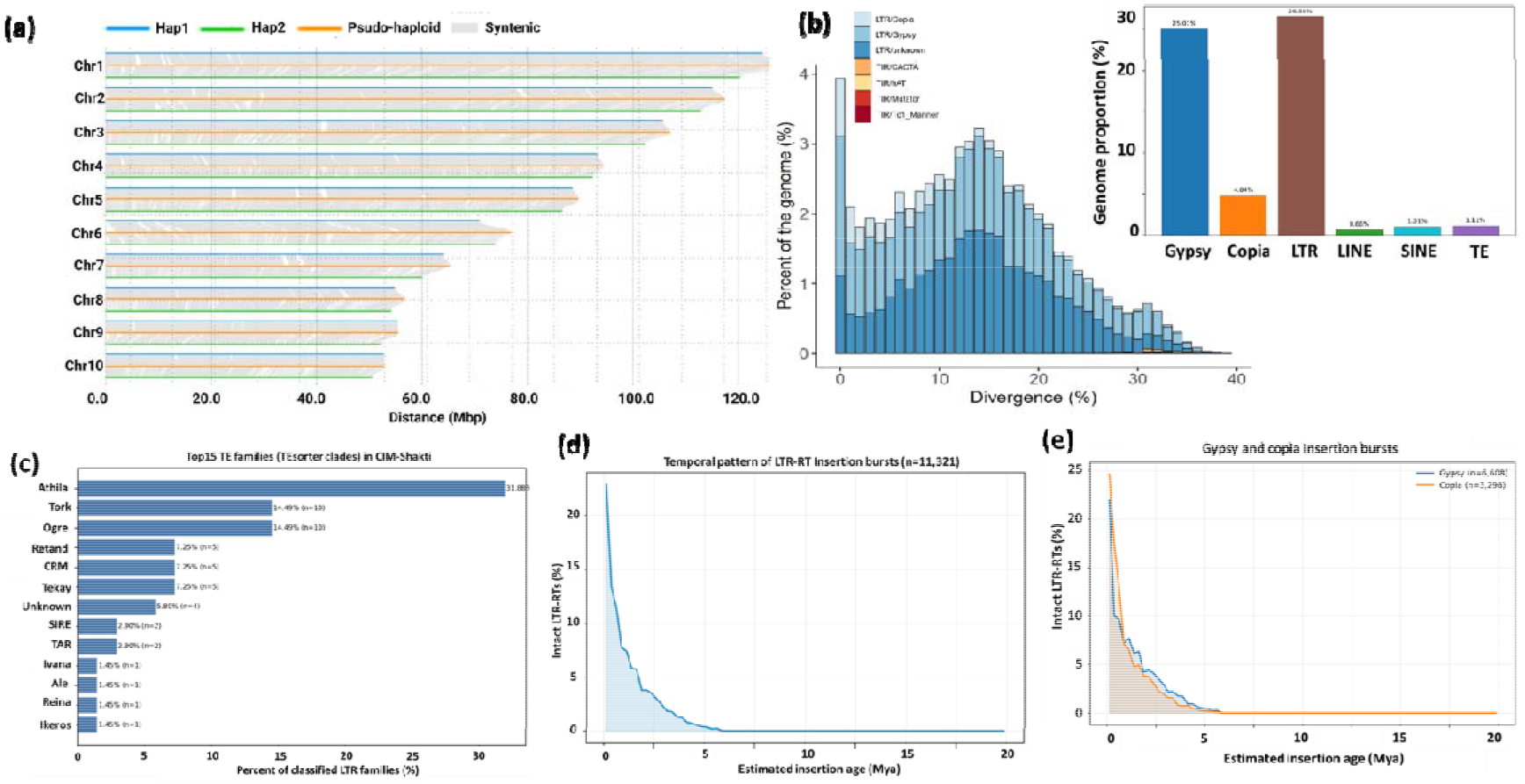
Comparative genome structure and repeat landscape of *Asparagus racemosus* var. CIM-Shakti. **(a)** Comparative chromosome-scale synteny among haplotype-resolved assemblies (Hap1 and Hap2) and the pseudo-haploid reference assembly of *A. racemosus* var. CIM-Shakti. Colored horizontal bars represent chromosome lengths, while grey connecting lines indicate conserved syntenic relationships across homologous chromosomes. **(b)** Repeat divergence landscape showing the genomic proportion and sequence divergence of major transposable element (TE) classes. LTR retrotransposons, particularly Gypsy and Copia elements, constitute the dominant repeat fractions of the genome. **(c)** Genome-wide distribution of major repeat classes. LTR retrotransposons were the predominant repetitive components, with Gypsy elements accounting for 25.01% of the genome, followed by unclassified LTR elements (26.58%) and Copia elements (4.84%), whereas LINEs, SINEs, and DNA transposons contributed relatively smaller proportions. **(d)** Distribution of the top 15 classified LTR retrotransposon families identified in the CIM-Shakti genome. Athila was the most abundant family, followed by Tork and Ogre lineages, indicating substantial expansion of Gypsy-related elements. **(e)** Temporal dynamics of intact LTR retrotransposon insertion bursts inferred from sequence divergence analysis. Most LTR-retrotransposon insertions occurred within the recent evolutionary past, with a major burst concentrated within the last 2–5 million years. **(f)** Comparative insertion-age profiles of Gypsy and Copia LTR retrotransposons showing recent amplification activity, particularly for Gypsy elements, suggesting their major contribution to genome expansion in *A. racemosus*.

### Pseudo-haploid assembly repeat annotation

Transposable elements (TEs) in the *CIM-shakti* genome was identified and annotated using an integrated pipeline that combined TRF (v.4.10), RECON, RepeatScout (v. 1.0.7), RepeatMasker (v4.1. 9), RepeatAfterMe (v.0.0.7)^15^. *De novo* repeat libraries were constructed with RepeatModeler (v2.0.7) ^16^ and EDTA (v2.2.0) ^17^ to improve classification accuracy. In total, 51,743,703 interspersed repeat elements were annotated, collectively representing 71.06% of the genome (Fig. 2b). These comprised both retrotransposons—long terminal repeat (LTR) and non-LTR elements—and DNA transposons with terminal inverted repeats (TIR) and non-TIR structures (Fig. 2b-e). Among these, LTR-Gypsy retrotransposons were the predominant class, accounting for more than 29% of the assembled genome (Fig. 2b-e).

### Pseudo-haploid assembly gene prediction and annotation

Gene prediction in the *CIM shkati* genome was performed using the MAKER pipeline (v3.01.03)^18^ which integrates ab initio predictions, protein homology, and transcriptomic evidence. Ab initio gene models were generated with Augustus (v3.4.0) ^19^, trained with *A. officinalis*. For transcriptome-based support, RNA-seq datasets derived from *A. racemosus* leaves, stem, root and inflorescences (public genome submitted to NCBI) were aligned with HISAT2 (v2.2.1)^20^ and assembled using StringTie (v2.1.7)^21^. Homology-based annotations incorporated protein sequences from *A. setaceus* and *Arabidopsis thaliana*. The integrated pipeline predicted a total of 29,199 protein-coding genes.

Functional annotation was carried out using a combination of approaches. Predicted proteins were searched against SwissProt/UniProt, NCBI nr, and Pfam (v35.0) using BLASTP and HMMER (v3.3.2). Functional domains were further characterized with InterProScan (v5.56-89.0) ^22^, and and Gene Ontology (GO) terms were assigned using Blast2GO (v6.0). Orthology-based annotation with eggNOG-mapper (v5.0)^23^ provided additional functional assignments, while the KEGG Automatic Annotation Server^24^ enabled mapping of genes to metabolic and signalling pathways. Functional annotation of the 29,199 predicted genes identified 16,768 genes with eggNOG-mapper and seed ortholog matches, 15,407 genes with Pfam domains, and 8,671 and 8,783 genes associated with GO and KEGG terms, respectively (Table 5). Genes involved in steroidal saponin biosynthesis pathways were specifically identified and manually curated based on homology to characterized genes recovered from the uniprot database.

## Results

### Genome assembly quality and structural validation

The chromosome-scale pseudo-haploid reference assembly of *Asparagus racemosus* (‘CIM-Shakti’) exhibited high contiguity, completeness, and structural integrity, providing a robust representation of the genome. The assembly spanned 817.5 Mb with a scaffold N50 of 98.50 Mb, indicating that a substantial proportion of the genome is contained within long, well-resolved chromosomes. In comparison, the haplotype-resolved assemblies (hap1 and hap2) showed slightly reduced contiguity, reflected by lower N50 values and increased fragmentation, which is expected due to the retention of allelic variation and incomplete phasing across complex or repetitive regions. In addition to superior contiguity, the reference assembly exhibited the lowest gap burden among the three assemblies (0.00784%), with fewer unresolved regions and minimal stretches of ambiguous bases. This reduced gap fraction suggests more effective resolution of repetitive and structurally complex genomic regions, likely facilitated by high-fidelity long-read sequencing and optimized scaffolding strategies. Collectively, these metrics indicate that the pseudo-haploid assembly achieves an optimal balance between completeness and redundancy reduction, making it the most suitable assembly for downstream analyses, including gene annotation, comparative genomics, and structural variation assessment. Gene space completeness was strongly supported by BUSCO analysis, Across the Eukaryota dataset, all three assemblies exhibited near-perfect completeness (99.8%), with 95.3% single-copy BUSCOs and only 4.5% duplicated genes. Fragmented BUSCOs were negligible (0.2%), and no missing BUSCOs were detected, indicating highly complete core gene recovery across all assemblies. For the Embryophyta dataset, completeness remained high (∼98.0–98.1%) across assemblies. The pseudo-haploid assembly showed the highest proportion of single-copy BUSCOs (94.0%) and the lowest duplication level (4.0%), compared to hap1 (4.9%) and hap2 (4.8%), further supporting its suitability as a reference genome. Fragmented (0.5–0.7%) and missing BUSCOs (1.2– 1.5%) remained low across all assemblies. In the more lineage-specific liliopsida dataset, completeness ranged from 97.6% in the pseudo-haploid assembly to 98.4% in hap2. Haplotype assemblies exhibited slightly higher duplication levels (∼4.0–4.1%) compared to the pseudo-haploid assembly (2.7%), consistent with retained heterozygosity. The pseudo-haploid assembly showed a modest increase in missing BUSCOs (3.4%), reflecting the expected trade-off between redundancy removal and completeness. BlobToolKit profiling of the CIM-Shakti genome assembly dataset included 53 scaffolds spanning 817567924 bp.

### Chromosome architecture and genome organization

Analysis of chromosomal features across the first 10 chromosomes revealed hallmarks of near-complete genome assembly. Canonical plant telomeric repeats (TTTAGGG) were detected at both ends of all chromosomes, indicating successful recovery of chromosome termini. Centromeric regions were consistently identified across all chromosomes, with an average of ∼100 tandem repeat–based candidate regions per chromosome. The highest-confidence centromeric intervals showed an average span of ∼267 kb and ∼10.4% tandem repeat coverage, consistent with typical plant centromere organization.

Genome-wide GC content analysis revealed a relatively uniform distribution, with a mean GC content of 40.2% across 1 Mb windows. Integration of GC content with telomeric and centromeric annotations provided a coherent chromosome-scale view of genome organization.

### Repeat annotation

Transposable elements in the Asparagus racemosus var. CIM-Shakti pseudo-haploid assembly were annotated using EDTA. Across 53 scaffolds (assembly span: 817,567, 924bp), 580,929,729 bp were masked as interspersed repeats, corresponding to 59.21% of the genome. LTR retrotransposons were the dominant component, with Gypsy elements masking 245,426,141 bp (25.01%), Copia elements 47,457,599 bp (4.84%), and unclassified LTR elements 260,741,047 bp (26.58%). Additional repeat classes included SINE-unknown (9,942,683 bp; 1.01%), LINE-L1 (4,260,132 bp; 0.43%), LINE-RTE (2,185,128 bp; 0.22%), CACTA (1,873,302 bp; 0.19%), and repeat fragments (7,329,843 bp; 0.75%), while other DNA transposon and helitron classes each contributed <0.1%. These results indicate that the CIM-Shakti genome is strongly enriched for LTR retrotransposons, especially Gypsy-related and unclassified LTR fractions.

### Gene structure and annotation landscape

The final structural annotation of the *Asparagus racemosus* var. CIM-Shakti pseudo-haploid genome identified 29,199 protein-coding genes and an equal number of transcript models, indicating that one representative transcript was retained per gene for the reference annotation. These gene models comprised 138,433 exons and 125,201 coding sequence features. The average gene length was 4,605.1 bp, with an average of 4.74 exons and 4.29 CDS features per transcript. Only 642 transcripts, representing 2.20% of the annotation set, were single-exon models, whereas 28,557 transcripts, representing 97.80%, contained multiple exons. This high proportion of multi-exon gene models supports the quality of the structural annotation and suggests effective integration of transcript, homology, and ab initio evidence during gene prediction. The recovery of a compact but structurally rich gene set is consistent with a high-quality chromosome-scale medicinal plant genome. The predominance of multi-exon genes also indicates that the annotation is not dominated by fragmented or short ab initio predictions, which can occur in repeat-rich plant genomes. Together with the high BUSCO completeness of the genome assembly, these gene structure metrics support the suitability of the CIM-Shakti reference for downstream functional genomics, pathway reconstruction, comparative genomics, and candidate gene discovery related to steroidal saponin biosynthesis.

### Data Records

The complete genome assembly and associated datasets have been deposited in public repositories with the given accessions. The final chromosome-level genome assembly of *A. racemosus* var. *CIM-Shakti* has been deposited at inhouse data repository and will be available after the manuscript publication.

### Technical Validation

The completeness of the gene space in the *CIM-Shakti* genome assembly was evaluated using BUSCO (v5.4.6) (36) with the *embryophyta_odb10* dataset (n = 1,614). The analysis recovered 98.6% complete genes, comprising 92.7% single-copy and 5.9% duplicated, with 1.0% fragmented and 0.4% missing genes. These results are on par with other high-quality grass genome assemblies, underscoring the comprehensive recovery of the coding repertoire.

## Code Availability

All computational analyses were performed using publicly available software packages with version numbers and parameters specified in the Methods section. Custom scripts for data processing, analysis pipelines, and visualization were developed using Python 3.8, R 4.2.0, and Bash scripting. No proprietary software was used in this study. All genome assembly and annotation pipelines can be reproduced using the provided documentation and publicly available computational resources. Specific software versions and database versions used are documented in the GitHub repository for complete reproducibility.

## Acknowledgements

We acknowledge the support of the Director, CSIR–Central Institute of Medicinal and Aromatic Plants (CSIR-CIMAP), under grant MLP002610.

